# 2-photon laser printing to mechanically stimulate multicellular systems in 3D

**DOI:** 10.1101/2023.12.23.573049

**Authors:** Federico Colombo, Mohammadreza Taale, Fereydoon Taheri, Maria Villiou, Teresa Debatin, Gent Dulatahu, Philipp Kollenz, Målin Schmidt, Christina Schlagheck, Joachim Wittbrodt, Christine Selhuber-Unkel

## Abstract

Most biological activities take place in 3D environments, where cells communicate with each other in various directions and are located in a defined, often microstructured, space. To investigate the effect of defined cyclic mechanical forces on a multicellular system, we develop a sub-millimeter sized stretching device for mechanical stimulation of a structurally restricted, soft multicellular microenvironment. For the stretching device, a multimaterial 3D microstructure made of PDMS and gelatine-based hydrogel is printed via 2-photon polymerization (2PP) method. The printed structures are first characterized microscopically and mechanically to study the effect of different printing parameters. With 2PP, organotypic cell cultures are then directly printed into the hydrogel structures to achieve true 3D cell culture systems. These are mechanically stimulated with a cantilever by indenting the stretching device at a defined point. As a most important result, the cells in the 3D organotypic cell culture change morphology and actin orientation when exposed to cyclic mechanical stretch, even within short timescales of just 30 minutes. As a proof of concept, we encapsulated a Medaka retinal organoid in the same structure to demonstrate that even preformed organoids can be stimulated by our method. The results demonstrate the power of 2PP to manufacturing multifunctional soft devices for mechanically controlling multicellular systems at micrometer resolution and thus mimicking mechanical stress situations, as they occur *in vivo*.

## 1. Introduction

Forces are undoubtedly essential factors in life and are continuously applied in all living systems, particularly to cells and their multicellular arrangements, where they can control development and morphogenesis[1] and cell metabolism[2]. Different compartments of cells have been identified as sensors of mechanical force, including the nucleus[3] and focal adhesion clusters[4]. As main mediators of force transmission, proteins such as vinculin, talin, paxillin and focal adhesion kinase have been identified[5–8].

A lot of progress has been made in recent years to investigate the effect of mechanical stretching on single cells[9,10] and on multicellular assemblies[11,12], including 3D stretching[13,14]. Typically, such forces are applied to cell sheets on or in elastic materials, which lack both micrometer precision and structural control of cell positioning at the micrometer scale, which would be highly valuable to mimic specific human tissue environments. For example, pathogenic conditions like osteoarthritis and rheumatoid arthritis affect the joints and are well known to be driven by the interplay of different cell types such fibroblast-like synoviocytes, macrophages and endothelial cells[15]. In this context, the formation of new blood vessels (angiogenesis), the thickening of the fibroblasts layers and the spatial distribution of the cells into the tissue have a key role in the disease development[16]. In these processes, the activation of pro- or anti-inflammatory pathways are also dependent on the mechanical forces perceived by the cells. Indeed, it is known that fibroblast-like synoviocytes are sensitive to shear stress[17] and to mechanical stretch[18,19]. However, these previous works have limitations, as they investigated the response to mechanical stimuli by maintaining cells in a traditional 2D cell culture. Hence, there is a need to provide methods to study the effects of mechanical stimuli at a higher level of complexity, maintaining cells in a 3D microenvironment under well-defined stress, and with multiple cell types simultaneously.

Using two-photon polymerization (2PP) 3D printing it is now possible to build high-resolution structures[20] at the micro- and nanometer scale for biomedical applications[21,22]. For example, complex structures have been designed and printed for studying the interaction of neurons [23] and in combination with responsive molecules 2PP-printed microstructures can even be controlled by external stimuli[24,25]. The possibility of printing such sophisticated structures with micrometer resolution makes it possible to study the impact of forces on individual cells. In this way, the forces produced by single beating cardiomyocytes could be observed by monitoring the deformation of microprinted structures[26] and inversely, a responsive structure based on host-guest reactions in a polymer was employed to mechanically stimulate individual cells and investigate the cellular response to such a stimulus[27]. Also, photoresponsive molecules have been proposed to induce mechanostimulation of cell adhesion and the upregulation of proteins associated with focal adhesion clusters[28].

In spite of the fascinating results generated with such studies on single cells, most cells naturally occur in multicellular systems, thus it is an essential task to understand how a defined group of cells or how various cell types in co-culture react to external force stimuli, particularly when they are grown in a 3-dimensional environment[29,30]. Indeed, the combination of multiple cell types supported by an extracellular matrix (ECM) has recently become very popular because of the similarities with the original human tissue or organ[29]. The advantages of using these 3D cell cultures are tremendous because they enable to study of numerous biological aspects *in vitro* with a complexity that was not previously possible with normal 2D cultures using standard petri dishes or flasks[29].

In this work, we show the responses to forces, applied by a controlled, oscillatory asymmetric stretch to an organotypic cell culture. Due to advances in co-culturing endothelial cells and fibroblast-like synoviocytes together, the extracellular matrix has been shown to be sufficient to recreate a 3D synovial organotypic cell culture similar to the human synovial tissue[16] Therefore, we here focus on the oscillatory mechanical stimulation of such co-culture systems in a 3D cylindrical micro environment to force the cells to arrange themselves concentrically, thus mimicking synovial tissue, where endothelial cells of the blood vessels are surrounded by fibroblasts[31]. With this system we can resemble the force that is, for example, present in the knee joint. Hence, we show how to mechanically stimulate at different frequencies 3D cell cultures in a highly controlled manner, both structurally and regarding the applied stretch. Using this proposed device, we also performed a preliminary study to mechanically stimulate Medaka retinal organoid. Controlled mechanical stimulation of this model organoid, could elucidate in the future the role exerted by mechanical forces on the development of this organ and on understanding molecular mechanisms underlying frequent pathologies of the organ deputed to vision.

## 2. Results and discussion

### 2.1. Design of the device for the mechanical stimulation of 3D microstructures

We have developed a structure to mechanically stimulate multicellular systems at the micrometer scale by combining different inks printed using two-photon polymerization lithography. We used a commercial ink IP-PDMS (Polydimethylsiloxane) as an elastic material, and two commercial hydrogels (see Materials and Methods) which are compatible with 2PP method. By means of simple Finite element analysis (FEA)[32] performed with Autodesk Inventor, we were able to predict the strain performing the simulations after applying a force of 200 µN. This strategy allowed us to apply asymmetric stress to cells grown in or on the hydrogel, respectively. According to the simulations we would get a maximum lateral displacement of about 3 μm while experimentally our data showed lower values (2 μm). This discrepancy is due to the fact that the parameters relating to the physical properties of the materials used in the simulation were adapted from the literature and were not obtained experimentally (the parameters used were given in Materials and Methods). Although the FEA thus only gave us approximate measurements, it was very useful to optimize the 3D model of the stretching device before it was printed. Furthermore, from this preliminary analysis, we were able to confirm that the type of lateral displacement obtained with the device will be asymmetric, with a major stress point close to the hydrogel. This situation can mimic, for example, the stress applied to cells in the knee joint[18]. In **Figure 1a** we showed the rendering of the CAD design (top and lateral view) of the 3D model that we used for FEA analysis (**Figure 1b**). In **Figure 1c, d** we show a sketch of the cultured cells before (**Figure 1c**) and after (**Figure 1d**) applying the mechanical stimulation with a cantilever.

**Figure 1.**
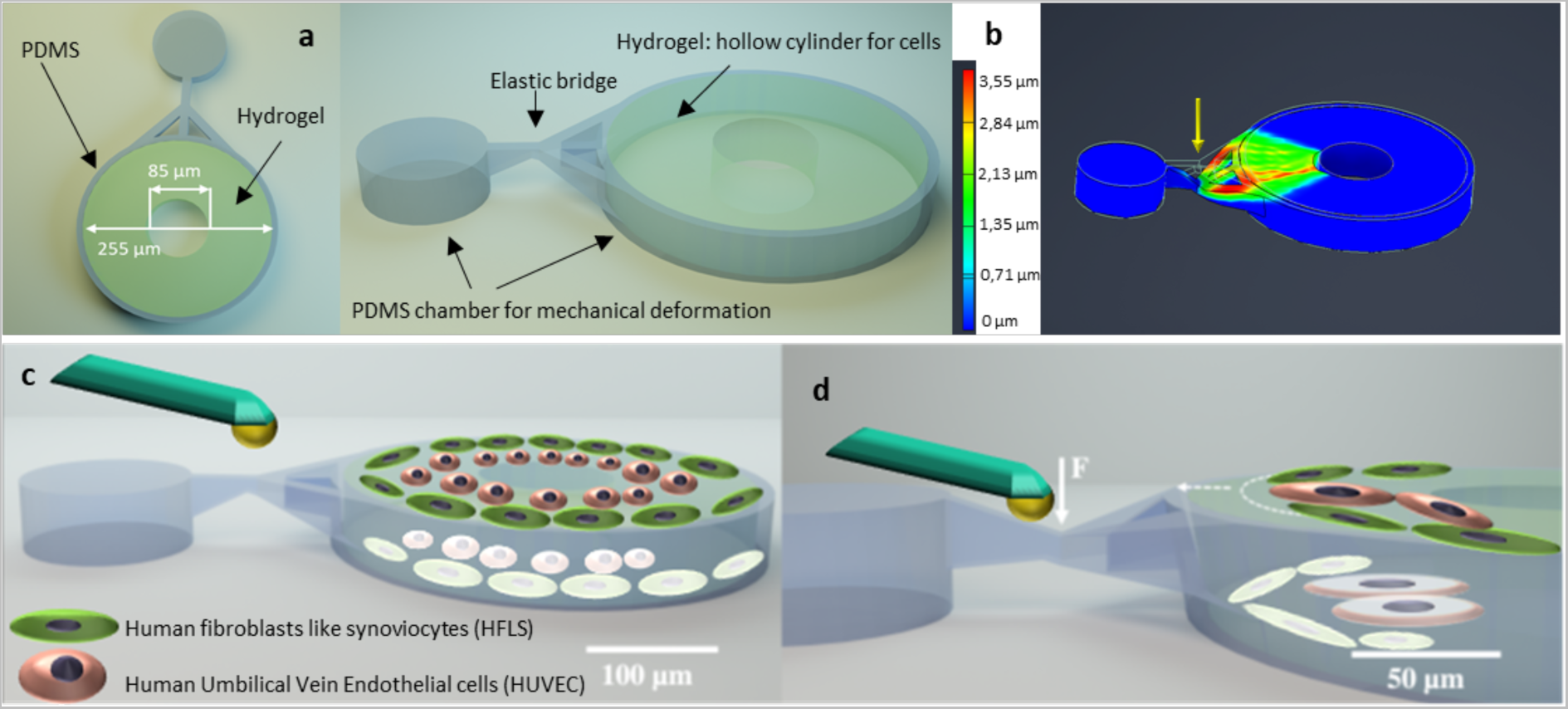
Overview of the 3D model used for the organotypic cell culture system. a) CAD model of the structure used for the prints. In light green the hollow cylinder made by hydrogel for the 3D cell culture system is shown. b) Results of the FEA analysis applying a load of 200 μN with a cantilever on the elastic bridge of the model. As shown an asymmetric and gradual displacement is obtained by indentation. c-d) Schematic drawing of the cells inside the 3D model and how they are mechanically stimulated.

### 2.2. Characterization of the hydrogel inks

To check the printing settings and mechanical properties of the hydrogel inks that we used to print the structures for growing cells, we printed an array of rings with a height of 10 μm, thickness of 10 μm and a total diameter of 100 μm using the ink HYDROBIO INX N100. The structures were printed using different combinations of laser power and scan speed. The lowest laser power used was 39.6 mW while the highest was 52.8 mW, instead for the speed we used a range from 45 mm/sec to 100 mm/sec. The quality of the print was assessed by bright field microscopy (**Figure 2a**), using different magnifications (10x, 20x, 40x). Since cells strongly react to the mechanical properties of their surroundings by changing adhesion and migration[33–35], we investigated the mechanical properties of the printed hydrogel materials as a function of laser power and printing speed by using a nanoindenter. We indented at least four points on the various rings printed with this ink. As shown in **Figure 2b**, the increase in laser power causes an increase in the stiffness of the structures. While the printing speed has less influence, the combination of the two clearly leads to an increase in stiffness. Despite these variations, the values we obtained vary from 250 Pa to 500 Pa on average, values that agree with what can be found in the literature for commercial extracellular matrices such as Matrigel[36]. Indeed, in the literature is known that the Young’s modulus increases with laser power [37,38]. In **Figure 2c** we showed one representative image acquired during the indentations performed on a ring printed with a laser power of 46.2 mW and speed of 100mm/s. The same characterization was performed on the HYDROBIO INX N400, which will be then used to print the 3D organotypic cell culture (**Figure S1**). In **Figure 2d-e** we showed cells grown on hydrogel cylinders that are printed with different laser power and thus different stiffness. After printing, the sample was incubated with a primary human fibroblast-like synoviocyte (HFLS) cell line, which is then used to print the 3D organotypic cell culture. We monitored the behavior of the cells in live cell imaging using the bright field microscope. After 1h and 30 minutes, the cells were already making stable contacts with the printed structures (**Figure 2d**). One or more cells were able to fill the inner part of the rings adopting the curvature of the structures, but we could not observe cells able to lay on the edges of the rings, probably due to the limited width of the wall ring structure (10 μm). Afterward, cells on the sample were labeled by immunofluorescence staining (**Figure. 2e**). Several cells were spread on the glass coverslip and few of the cells were inside the structures showing that they were adapting their shapes to the rings. DAPI was used to stain the nuclei, phalloidin was used to stain the actin and the anti-paxillin was then used to highlight the focal adhesions which were expressed both in cells in- and outside the rings. As shown in these images, the cells also adhere to the glass slide, as this has not been treated to prevent cell adhesion. The actin is homogeneously distributed within the various cells covering the slide and also within those making contact with the cylinders, while Paxillin staining shows the accumulations of the focal adhesion between the cells and the hydrogel (see arrow in the zoom-in **Figure 2e**).

**Figure 2.**
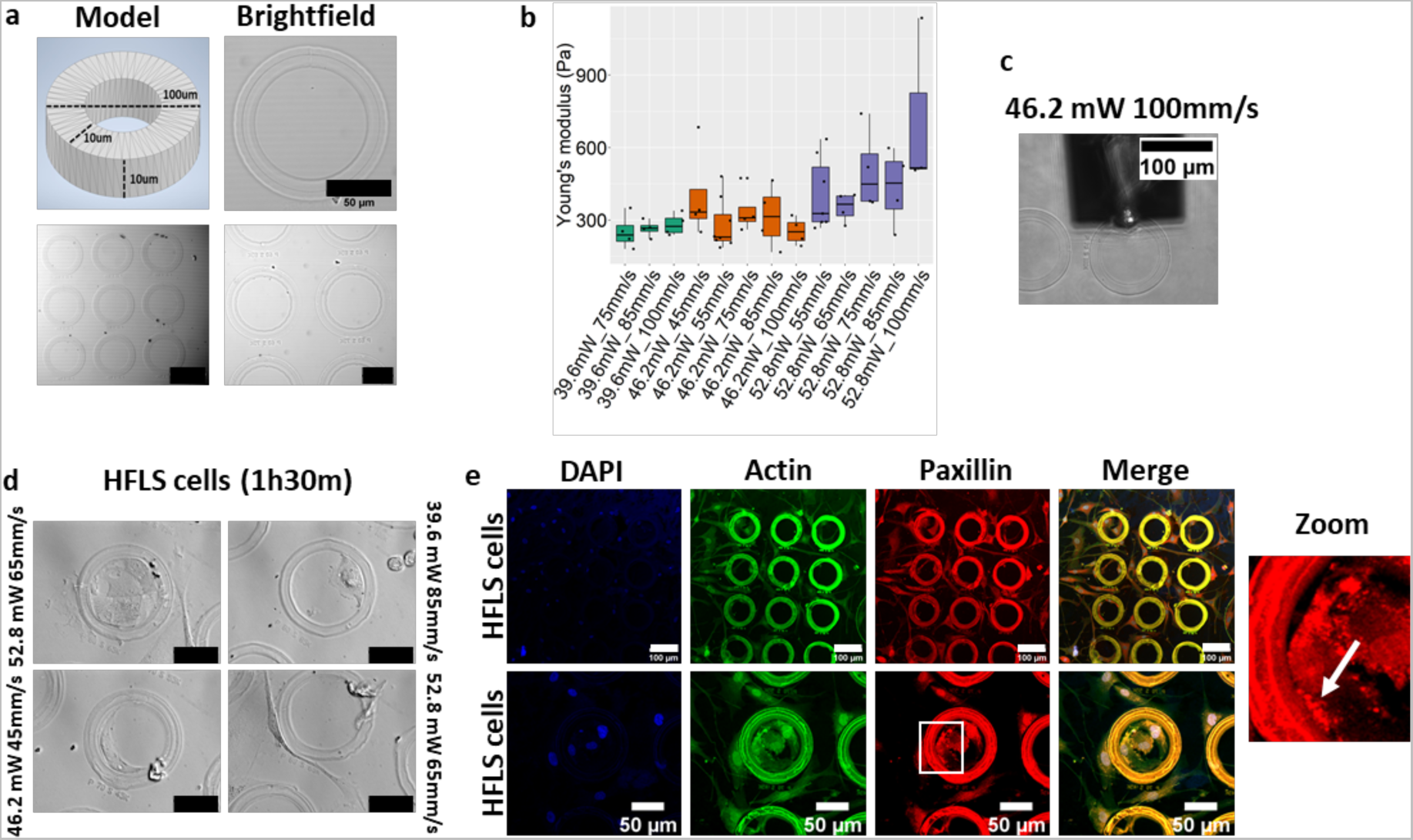
Characterization of the hydrogel ink. a) CAD model of the hollow pillar and brightfield images of them after the print. Scale bar of images on the right side is 50μm while on the left side is 100 μm. b) Boxplot of the Young’s Modulus (Pa) of different cylinders printed with HYDROBIO INX N100 ink with increasing laser power and speed. c) Representative image acquired during the analysis with the nanoindenter. The microscope coupled to the nanoindenter helped us drive the probe to the desired location. d) Brightfield images of the printed cylinders incubated for 90 minutes with HFLS cells. Scale bars 50μm. e) Fluorescence analysis on the cells incubated with the hydrogel cylinders. Cells were stained for Actin (Phalloidin-Alexa488 in green), for Paxillin (red) and nuclei (DAPI). All the structures presented in this figure belong to the same sample and are therefore printed using the same settings. Scale bars top row 100 μm while in the bottom row the scale bars are 50μm.

### 2.3. Characterization of the 3D structures after print

After testing the growth of the cells on the hydrogel HYDROBIO INX N100, we printed the previously designed 3D stimulation devices in their complete version made with a cylindrical hydrogel core (HYDROBIO INX N100) surrounded by an elastic chamber (IP-PDMS). As shown in **Figure 3a**, scanning electron microscope (SEM) images showed the high print quality of the external elastic chamber. We focused our attention on the bridge section because it will be the part where indentation and thus mechanical stimulation should take place. In addition, from the top view is possible to detect the two stitching points that cannot be avoided due to the size of the print (total length > 300μm). Despite these two stitches, the structure, as will be shown later during the stimuli, resists mechanical stimulations without being damaged. Then, using a fluorescence microscope we also verified the print quality of the entire 3D model by exploiting the autofluorescence of the IP-PDMS at 405nm and that of the hydrogel at 488nm (**Figure 3b**). In the boxplot in **Figure 3c**, the % of swelling/shrinkage of three printed models is shown. All the features of the structures measured show a difference of less than 10% compared with the CAD design file. Furthermore, we investigated the effects of printing the hydrogel cylinders with and without elastic structure in IP-PDMS. As demonstrated in **Figure S2**, the cylinders printed alone swell in comparison to those with IP-PDMS (with a diameter of 380 μm and 260 μm, respectively). To prove that the cells in general can grow onto the hydrogels of these devices, these were incubated with REF (Rat Embryonic Fibroblasts) cells for 4 days and finally stained using DAPI for nuclei, Phalloidin Alexa488nm for actin and anti-paxillin for focal adhesions (**Figure S3a-b**). We also observe that cells can adhere to PDMS structures as can be seen from **Figure S3 a-b**. This is most likely due to the fact that during the second printing step, part of the hydrogel is not eliminated and traces of this material persist as a coating of the structures. Moreover, the cells were shaping themselves according to the 3D hydrogel cylinder. This finding is very important because we can decide *a priori* the shape we would like to have in our 3D cell culture. This suggests that it would be possible to recreate desired cellular organizations and then study their spatial role in an organotypic cell culture with other cell lines, or to study their response to different mechanical or chemical stimuli.

**Figure 3.**
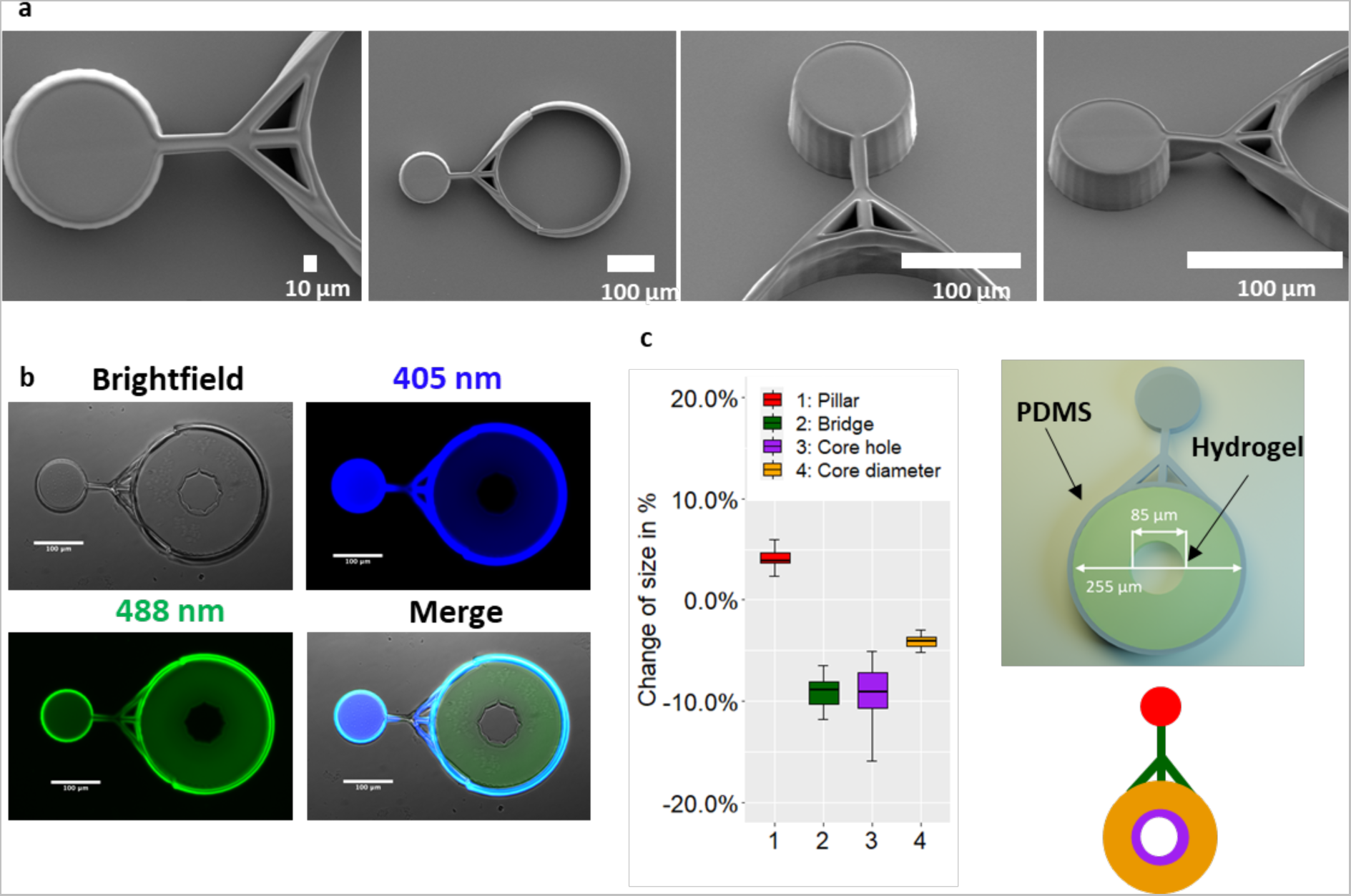
Characterization of the stretching device. a) SEM images of the IP-PDMS structure after print. b) Images of the final structures acquired at the fluorescence microscope. The IP-PDMS was autofluorescent at 405nm and the core hydrogel at 488nm. c) Boxplot of the swelling/shrinkage measured on the images taken at the fluorescent microscope. For each feature 6 different 3D structures were analyzed and data plotted as size variation in % from the size of the CAD file. On the right-side sketch of the features measured.

### 2.4. *In situ* printing of the cells

In the previous sections, we have shown how the 3D structures can be used to shape incubated cells on top of the structures (**Figure S3**). These are able to adhere to the hydrogel and infiltrate it. However, the next step is to directly print the cells within the hydrogel. Here, the circular shape of the hydrogel aims to recreate the inner ring of cells of an endothelial vessel. As proof of concept, HUVEC cells were cultured together with HFLS (human fibroblasts-like synoviocites). These two cell lines are known to self-assemble with the HUVEC cells forming the inner part of the vessel while the fibroblasts will form the underlying supporting tissue[16]. To achieve this, we used a commercial HYDROBIO INX N400 which is optimized for this purpose. To prove that this was possible, we directly printed the HFLS and HUVEC cells previously labelled respectively with a green and red dye. The method used to obtain this 3D organotypic cell culture is schematically represented in **Figure 4a**. During the *in-situ* cell printing process, we recorded a video (**Video S1** in the Supplementary Videos section) showing the shift in the focal plane as the hydrogel is formed, encapsulating the cells within. Figure S4 shows a frame image from the video before the printing started and another frame after the end of the printing process. The structures containing the organotypic cell culture were kept in culture for 5 days and then observed with a confocal microscope. As can be seen from Figure 4, the structures encapsulate a sufficient quantity of cells, with a ratio similar to that reported in the literature to form vascularized organoids, maintaining a ratio between 20% and 50% of the whole cell population[16,39]. In order to assess the viability of the cells after 2PP and to evaluate the toxicity of the hydrogel, we performed staining with calcein AM and propidium iodide (PI). **Figure S5** shows that most of the cells are alive (in green) while a few of them inside the structure were dead (positive for PI).

**Figure 4.**
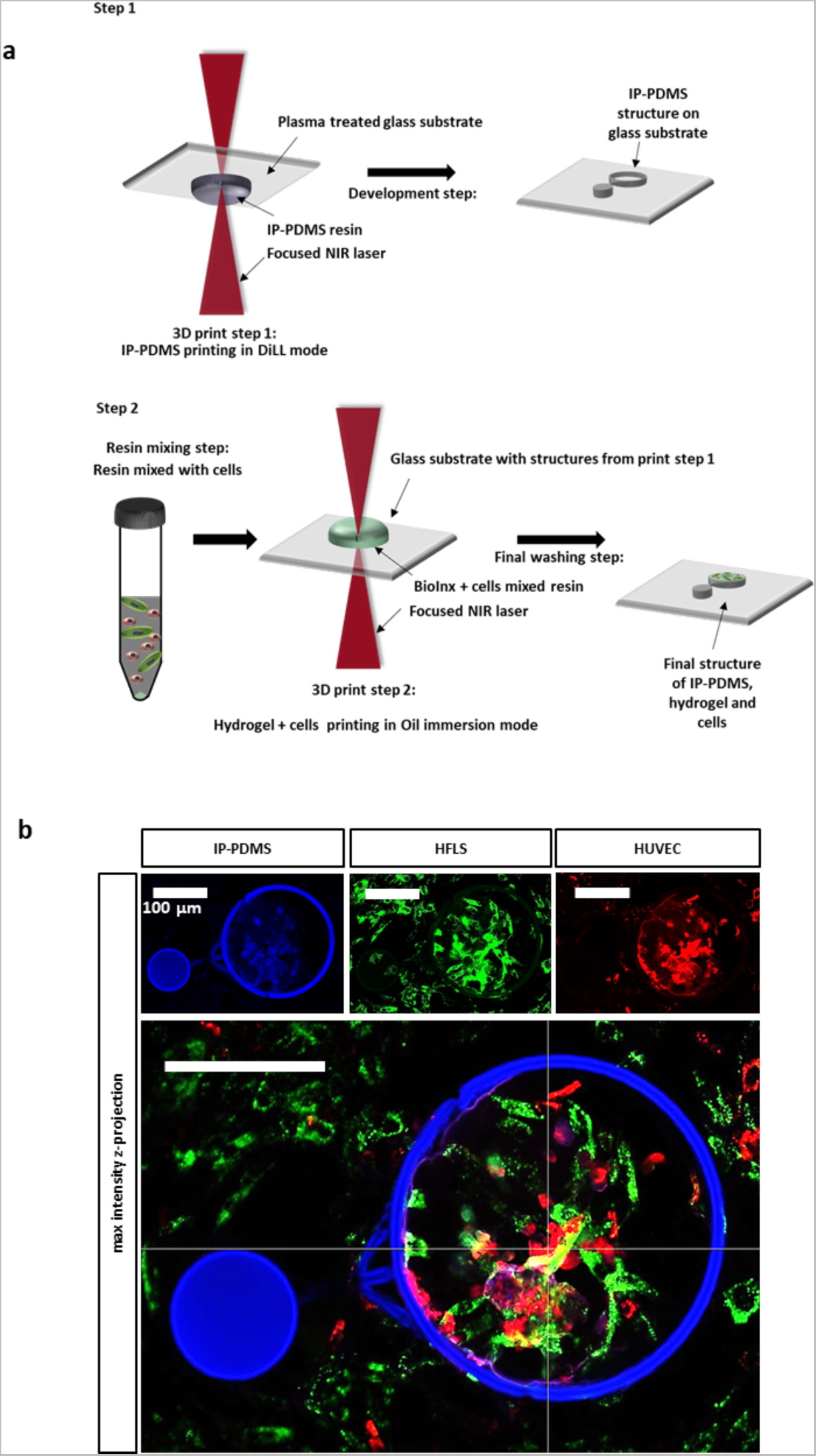
*In situ* 3D printing of the cells in the stretching device. a) Schematic presentation of the method used for the 2PP to print the cells in the hydrogel and generate the organotypic cell cultures. In the top row, the first step is to print the IP-PDMS structure. On the second row, the second step is where the cells are mixed with the resin to be directly printed in the hydrogel. b) Confocal images of the final structure printed with HFLS previously labeled with a green dye and HUVEC cells previously labeled with a red dye. In blue we acquired the autofluorescence of the IP-PDMS structure. Scale bars 100 μm.

### 2.5. Cantilever-driven mechanical deformation of the 3D printed organotypic cell culture systems

Most of the published work on mechanical stimulation performed on structures using the 2PP technique has focused on the cellular response to mechanical stimuli applied to a single-cell level[40]. However, the importance of studying cellular responses in more complex multicellular assemblies is well-known[30]. We therefore recreated a structure that could accommodate two mature cell lines as an organotypic cell culture system to study both their self-assembly and response to these stimuli. To control the mechanical stimulus, we used a nanoindenter equipped with a cantilever with a spring constant of 4.10 N/m and a sphere with a diameter of 24um. We utilized the cantilever to apply a precise force on the IP-PDMS bridge, inducing controlled displacement within the hydrogel cylinder containing the cells. In **Figure 5a** we showed brightfield images of a sample before and after 30 minutes of stimulation at 0.5 Hz (top row), and 1 Hz (middle row) using a load of 200 μN. The choice of stimulation frequency was based on previous studies that showed that 15 minutes of stimulation at a frequency of 0.5 Hz was sufficient to observe remodeling of the cytoskeleton[41]. The control sample was not stimulated but processed using the same conditions of the stimulated sample. In **Figure 5b** we analyzed the video recorded during the stimulations to generate the displacement map (See **Video S2** in the **Supplementary information**). To calculate the displacement we used the Lucas–Kanade optical flow method[42], and the data suggested that we can obtain a max displacement of 2μm in the area close to the stimulation for the samples stimulated at 0.5 Hz. In contrast, for samples stimulated with 1 Hz, we detected a lower displacement. This is due to the fact that hydrogels, as biological samples, behave as a viscoelastic material, consequently with increasing strain rate, the material becomes stiffer and stronger[43]. However, since the displacement is on purpose asymmetric, we analyzed the stimulations performed on the structures and we plotted the averaged displacement value for the first 100 frames (equal to 4.16 minutes) (**Figure 5c**). As shown, the average displacement value for the samples stimulated at 0.5 Hz is 0.25 μm, a displacement value which is in line with the displacement showed in previous work[27]. Whereas for structures stimulated at 1 Hz, the average displacement drops to 0.08 μm. The main advantage and novelty of our 3D structures lie in the possibility to incorporate cells that experience higher displacement in the proximity of the stimulation point, alongside cells that receive lesser or no stimulation. This will minimize variations between organotypic cell culture systems increasing the consistency of the data generated.

**Figure 5.**
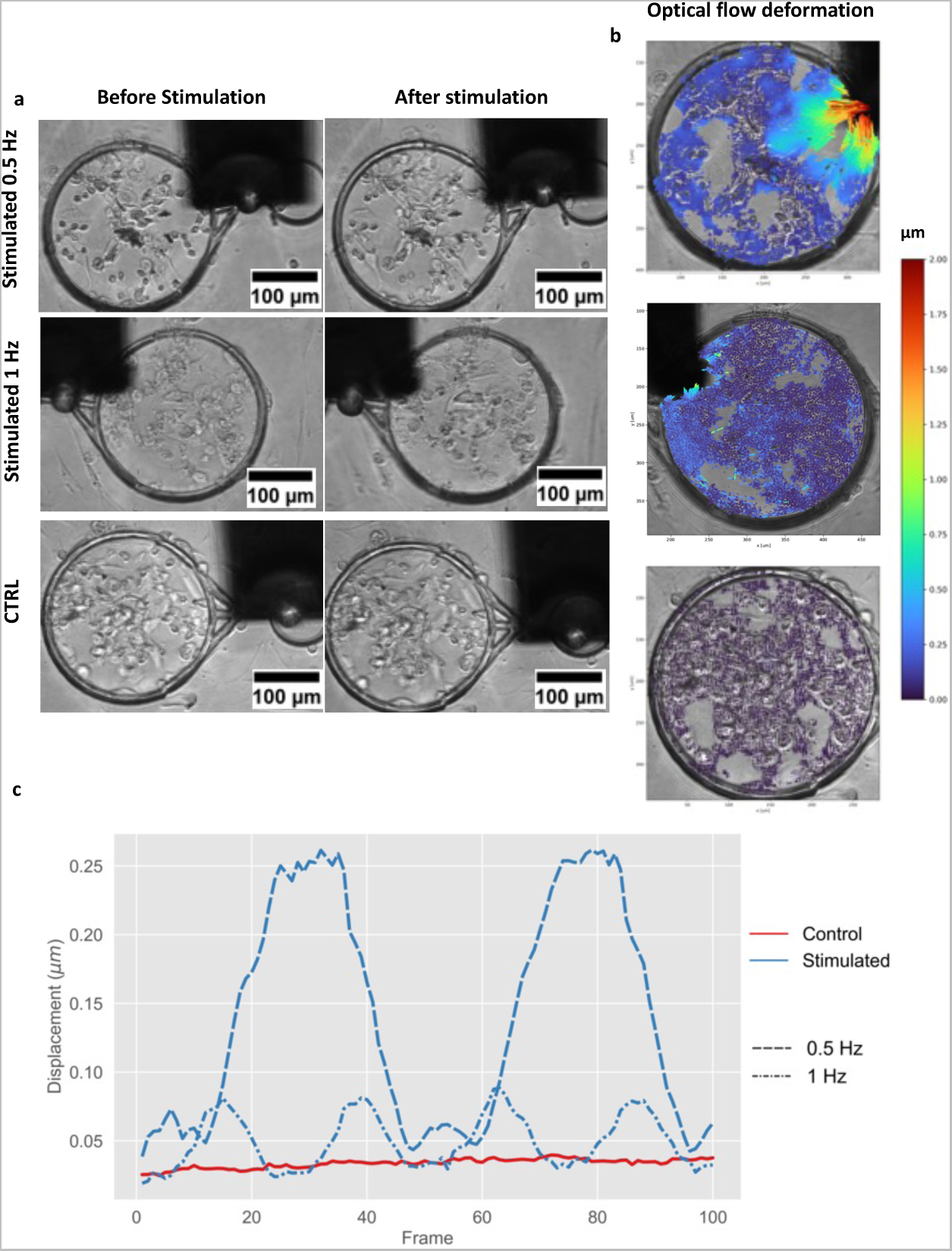
Mechanical stimulations of the organotypic cell culture systems. a) Brightfield images acquired before and after the mechanical stimulations (top row: 0.5 Hz; middle row: 1 Hz) and non-stimulated samples (bottom row). b) Displacement maps of the hydrogel with the cells (HFLS + HUVEC) were obtained using the optical flow method of stimulated samples and non-stimulated sample. For each condition we analyzed 3 samples. c) The graph shows the averaged displacement value of a structure stimulated at 0.5 Hz, a structure stimulated at 1 Hz and the control.

### 2.6. Mechanical stimulations at 0.5 Hz led to morphological changes in cells and their actin remodeling

After the mechanical stimulation the structures were placed in 4% PFA in order to fix the system and later perform immunofluorescence analysis. This will be used to understand the effects of mechanical stimulation on the organotypic cell culture systems. In **Figure 6a-b**, confocal images are shown respectively from 3 structures that were stimulated at 0.5 Hz and 3 structures stimulated at 1Hz, while in Figure 6c control images of 5 non-stimulated structures are shown. In **Figure 6** we show only one sample for each condition (see **Figure S6** for all the samples). In order to follow the self-assembly of the cells in response to the stretch, they were fluorescently marked before being mixed with the resin and printed. HFLS cells were marked in green while HUVEC cells were marked in red (in Figure 6 were falsely coloured in yellow). For each sample in the upper part we show the maximum intensity of different channels so that we can distinguish the HFLS from the HUVEC cells. In the bottom part the images of the actin and the results of the analysis performed to evaluate the actin alignment are presented. This analysis was performed with an algorithm recently published[44] which allows detecting the orientation angle of the aligned fibers, and the eccentricity which describes the elongation of the fibers in the region. A higher value of eccentricity means that we have longer fibers and a lower eccentricity means we have a more circular arrangement. Indeed, as shown in **Figure 6a**, we can observe that stimulated samples at 0.5 Hz have lower eccentricity suggesting that the cells have a more roundish morphology. Moreover, the cells (mainly HFLS in green) in the non-stimulated samples form a circular ring on the outer part of the structure while the HUVEC cells are arranged on the innermost part. In general, it is observed that the cells in the control group, as well as the cells in the structures stimulated at 1 Hz, are spread and obviously larger than the cells that were mechanically stimulated at 0.5 Hz. The latter, stimulated at 0.5 Hz, are evidently rounder and the actin staining shows few stress fibers compared to the control and 1 Hz stimulated structures, suggesting that this leads to actin remodeling. The presence of more rounded, shorter cells in samples subjected to higher strain is a phenomenon reported in previous studies[45]. All these evidences are supported by the histogram in **Figure 6d** which clearly shows the difference in the order parameter values analyzed in all the samples. The order parameter can take values between 1 and -1, where 1 is the complete alignment of the fibers, 0 is a random orientation, and -1 is the opposite alignment. By stimulating the structure at 0.5 Hz with higher displacement, we observed a significant disruption of the actin network hence a clear decay of order parameter is shown in **Figure 6d**. The effects of mechanical stretching have previously been studied both as an effect on single cell[27] and as effects on organoids[46] and 3D cell cultures using extracellular matrices[47]. These have been observed to lead to cell differentiation, cell proliferation and cell migration[48]. However, these studies show that the effects depend on the cell line studied, the duration of the mechanical stimulus and its frequency. During this process, the remodeling of the cytoskeleton copes with mechanical stress to prevent ruptures or cell damage[49]. Here, we therefore studied the effects caused on actin after a maximum asymmetric stretching of 2μm using a load of 200μN at 0.5Hz and a maximum asymmetric stretching of 0.5μm using a load of 200μN at 1 Hz. These findings show that, using this device, a higher stimulation frequency (e.g., 1 Hz) leads to lower displacement. This in turn has comparable effects to the control, with no appreciable difference in actin staining (**Figure 5b**). In contrast, a stimulation of 0.5 Hz caused actine remodeling and to understand other related biological impacts due to a lower frequency stimulation, clearly more detailed studies will be necessary.

**Figure 6.**
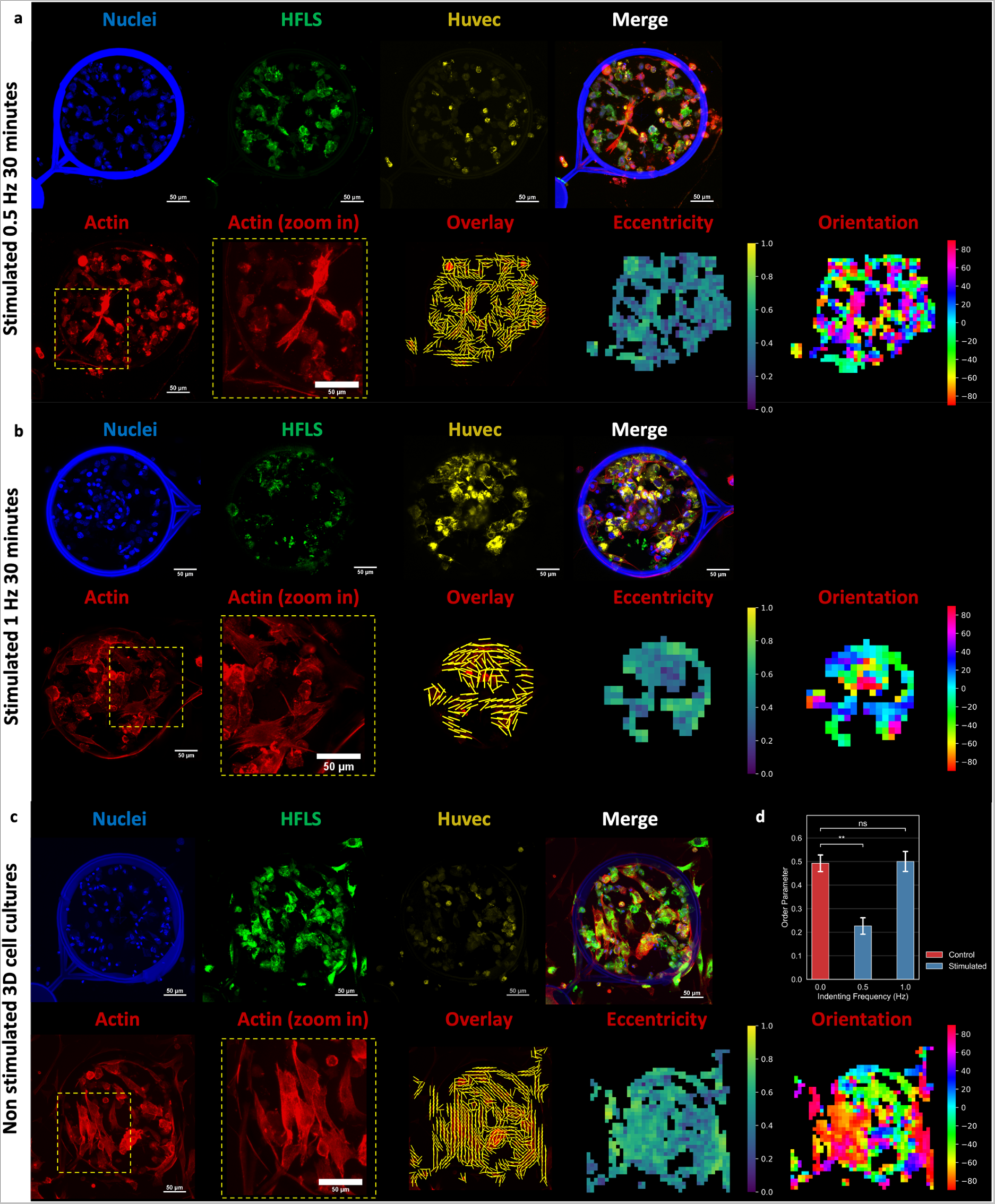
Morphological changes and actin remodeling of the mechanically stimulated organotypic cell culture systems. a) 3D organotypic cell culture systems (3 samples) mechanically stimulated for 30 minutes with a load of 200 μN at 0.5Hz. Here we present one sample for each stimulation or non-stimulation condition. Top row: max intensity projections of the 4 channels: HFLS cells (green), HUVEC cells (yellow), Nuclei (Hoechst) and merge. In the bottom row actin (red), zoom-in of the actin, and the results of the actin alignment as overlay, eccentricity and orientation angles. The overlay shows the alignment vectors in yellow while the FFT eccentricity map highlighted a circular orientation with a value close to 1 (yellow) and elongated with value close to 0 (purple). The orientation angles of actin fibers are displayed in the 2D orientation map. b) 3D organotypic cell culture systems (3 samples) mechanically stimulated for 30 minutes with a load of 200 μN at 1 Hz. Channel colors are the same as the stimulated samples at 0.5 Hz c) Control of the 3D organotypic cell culture systems non-stimulated. Channel colors are the same of the stimulated samples at 0.5 Hz. All the scale bars are 50μm. d) Histogram of the analysis performed on the actin alignment: 5 non-stimulated samples (red), 3 stimulated samples at 0.5 Hz, and 3 stimulated samples at 1 Hz. Statistical significance (p < 0.05) was tested by independent t-Test.

### 2.7. Mechanical stimulations of Medaka retinal organoid

As a proof of concept, in order to mechanically stimulate a preformed organoid as a more advanced 3D multicellular system, we encapsulated a Medaka retinal organoid in our device. Using this organ model based on fish primary embryonic pluripotent cells is beneficial because it quickly generates a system that closely resembles the human retina[50]. Additionally, its small size makes it an excellent candidate for 2PP printing, aligning with the aims of the proposed device. The procedure, requiring numerous steps and reported in **Figure 7**, was technically complicated overall. In particular, after placing the organoid inside the PDMS structure (**Figure 7a**), the critical moment was the exchange of the culture medium with the printing ink (hydrogel). Indeed, in this step the organoids undergo dehydration. This inevitably leads to the collapse of the organoid structure. Another challenge was the flow motion caused by the ink on the sample: this can lead to displacement of the organoid out of the structure. Nevertheless, we were able to obtain a good quality sample after printing (**Figure 7b**), which we stimulated using the same conditions used in the previous paragraph: a load of 200μN at 0.5Hz for 30 minutes. As shown in **Figure 7c**, obtained by confocal microscopy after stimulation, part of the organoid is encapsulated inside the hydrogel doughnut, while a portion is in its hole. The histones H2B are reported in green, while actin stained by SiR-Actin 652 nm is shown in red. Using the video captured during stimulation (see **Video S3** supplementary information), it was possible to calculate the average displacement (**Figure 7d**), which corresponds to about 0.20 μm. This is in line with the data collected for the displacement of the cellular structures stimulated in paragraph 2.5. These preliminary results, although obtained with a single sample, show that this device, despite the technical limitations related to the print, can be extremely useful for studying the effects of mechanical stretch on complex organoids such as that of the retina. The results could be extremely useful to understand the biological effects of mechanical stress during retinal development or to verify the possible effects of mechanical damage in pathologies.

**Figure 7.**
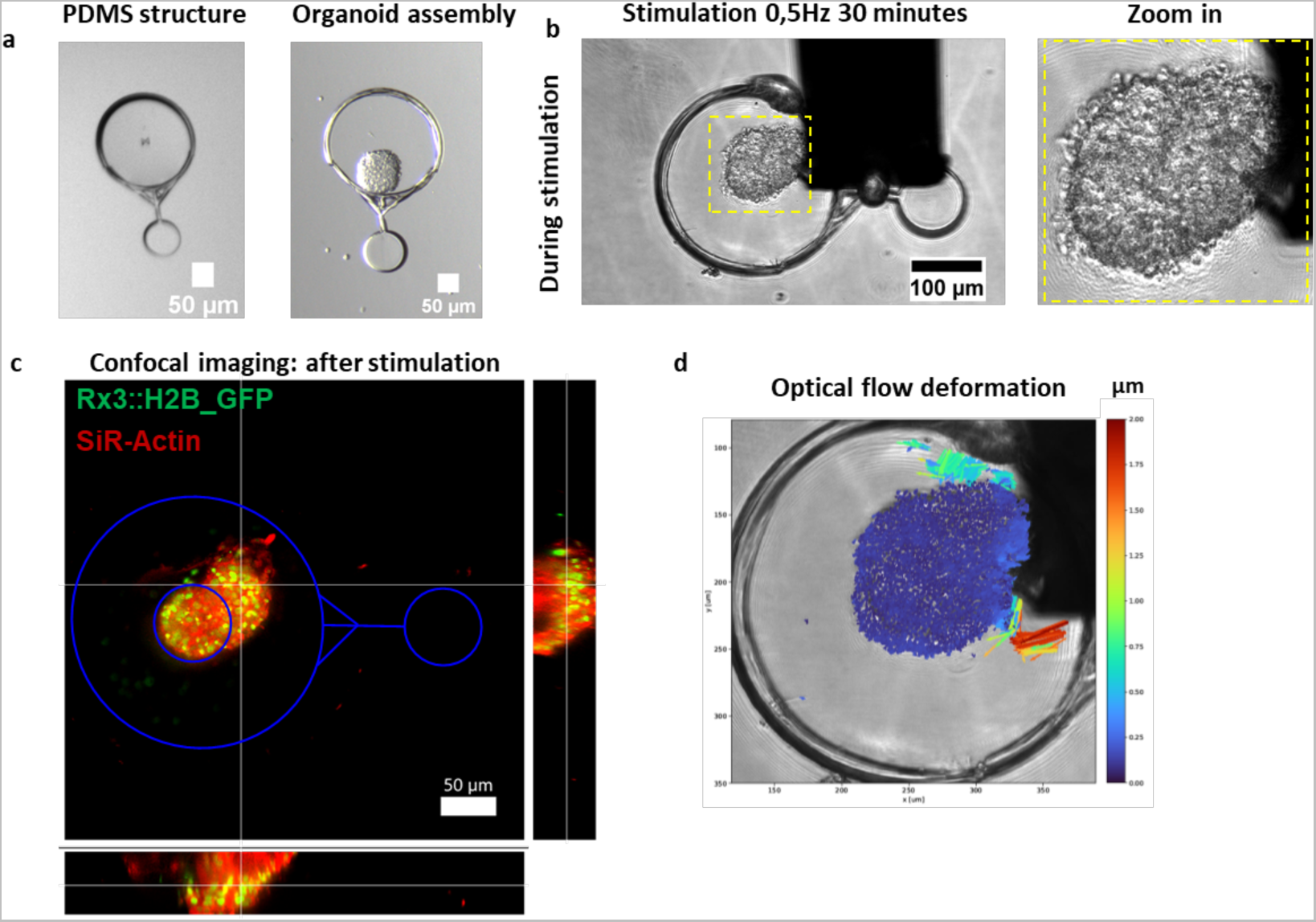
Mechanical stimulations of Medaka retinal organoid. a) Image of the PDMS structure before and after mounting the Medaka retinal organoid. b) Image of the complete structure with the organoid encapsulated within the hydrogel during the mechanical stimulation. An enlargement of the organoid is shown on the right. (c) Image obtained with a confocal microscope by acquiring the green fluorescence of the histons H2B and actin stained with SiR-actin at 652nm in red. In blue we drew the structure to facilitate the understanding of the positioning of the organoid inside the structure. (d) Displacement map obtained by the method shown in the paragraph 2.5. In this experiment, only one sample was analyzed as a proof of concept.

## 3. Conclusion

In this work, we have designed, fabricated and characterized a 3D microstructure device, printed via 2PP to generate a 3D organotypic cell culture that can be mechanically stimulated with unprecedented precision and controlled at the micrometer scale. The mechanical properties of the hydrogel inks were systematically investigated, revealing the influence of laser power and printing speed on the stiffness of the structures. The analysis of the 3D structures, made of a PDMS chamber and a hydrogel core, showed high print quality and minimal swelling/shrinkage compared to the CAD design, thanks to the structure design.

We printed two cell lines directly within the hydrogel creating a multicellular system. Mechanical stimulation of the organotypic cell culture was achieved by applying controlled forces using a cantilever-based device. A stimulation of 30 minutes at 0,5 Hz led to morphological changes and actin remodeling in the encapsulated cells, while a stimulation of 1 Hz was not effective due to less displacement into the structures. This emphasizes that increasing the frequency of stimulation for this structure and the material used (hydrogel-based) leads to less overall displacement. This device can be easily customized to obtain different degrees of stretch and to stimulate different types of multicellular systems both for short and long-term observations. Therefore, this work opens prospects in many biomedical applications, where mechanical stress situations help to elucidate functional features of multicellular systems, ranging from stem cell based organoid culture to the mechanical stimulation of cancer cell spheroids. Indeed, as a proof of concept, a Medaka retinal organoid was printed using the same device to demonstrate that this system can in the future be used to study the effects of mechanical stimulation on retinal development and to assess the impact of mechanical stress or damage in the pathogenesis of eye-related diseases.

## 4. Experimental Section/Methods

### 4.1. Two-Photon Polymerization

The 3D scaffold models used in this work were designed using Autodesk Inventor Professional 2023 and rendered with the open-source program Blender. Computer Aided Design (CAD) models (STL files) of the structures were loaded into the DeScribe 2.6 software (Nanoscribe GmbH). The printed 3D structures are based on two different material structures, a PDMS and a gelatin-based hydrogel. The PDMS structure forms the elastic backbone for force application. The hydrogel is the material where the cells are encapsulated. First, the elastic PDMS structure was printed onto a glass slide. Before applying the IP-PDMS resin (Nanoscribe GmbH) to the glass slide, its surface was activated by a 2-minute plasma treatment with a plasma pen (piezobrush®PZ3, relyon-plasma, Germany). The computer-aided design structure (the outer ring, bridge, and base) was printed at 25x, NA = 0.8 using a commercial laser direct writing device (Photonic Professional GT2, Nanoscribe GmbH). A laser power of 46.2 mW and a printing speed of 60 mm/sec was used as printing parameters. The structures were developed for 12 minutes using isopropanol (Sigma-Aldrich, Germany). To finalize the multimaterial structure, commercial gelatin-based hydrogel resin was used. For printing organotypic cell culture systems directly into the hydrogel the hydrogel resin was prepared by mixing 10 µl photoinitiator and 90 µl resin at 40 °C. The cell pellet (200.000 HFLS and 200.000 HUVEC cells, both pre-labeled before preparing the cells pellet) were immediately resuspended in the prepared resin and placed over the already printed PDMS structures. A 25X, NA = 0.8 oil immersion objective was used to print the cell-laden hydrogel within the PDMS structure. A laser power of 52.8 mW and a printing speed of 60 mm/sec were used as printing parameters for this part of the device. The structures were developed using the EBM-2 Basal Medium (CC-3156) supplemented with EGMTM-2 SingleQuotsTM Supplements (CC-4176) incubated at 37 °C. To prepare the print jobs, all models were hatched and sliced using DeScribe software (Nanoscribe GmbH).

To print the 3D structures that were simply incubated with cells on top we followed the same process described above without adding cells to the resin. In this case, the HYDROBIO INX N100 (Bioinx) was used.

### 4.2. Cell culture

The cell experiments were performed with three different cell lines. Rat embryonic fibroblast 52 wild type (REF52-wt, passage 10 - 25) were cultured with Gibco Dulbecco’s Modified Eagle Medium (DMEM) with 10% FBS and 1% Penicillin/Streptomycin (100 μg/mL). Human Umbilical Vein Endothelial Cells (HUVEC, passage 2-15) (Lonza, cat. Number C2519A) were cultured with EBM-2 Basal Medium (CC-3156) supplemented with EGMTM-2 SingleQuotsTM Supplements (CC-4176). Human Fibroblast-Like Synoviocyte (AddexBio, passage 3-15) were cultured with HFLS Growth Medium (CellApplications).

Cultivation and cell experiments were performed under sterile conditions. Cells were incubated with standard cell culture conditions at 37°C and 5% CO2. For cell splitting, the cells were washed twice with Phosphate-buffered saline (PBS), detached with 2-3mL of Trypsin for 5 minutes and centrifuged at 300 x g for 5 minutes.

### 4.3. Cell stainings

Prior to seeding the HFLS cells were labeled using the green CellMask™ Plasma Membrane staining kit (Invitrogen, cat. Number C37608) while the Huvec cells were labeled using the deep red CellMask™ Plasma Membrane staining kit (Invitrogen, cat. Number C10046) following the manufacturer’s protocol. After the cells were detached, pooled and centrifuged to obtain the cell pellet.

### 4.4. Staining of Paxillin, Actin and Nuclei

Cells growing on or inside the structures were fixed in 4% paraformaldehyde (PFA) for 1h. After the cells were washed three times with PBS for 5 minutes and blocked for 1h at room temperature (RT) with a blocking solution made of PBS and 10% fetal bovine serum (FBS) to reduce background during imaging. Then, the primary antibody rabbit anti-paxillin (Sigma-Aldrich cat. Number HPA051309) was diluted at a final concentration of 1 μg/mL in PBS and incubated overnight at 4°C. After one day, the samples were washed 3 times in PBS for 10 minutes each step. The secondary antibody goat anti-rabbit Alexa Fluor 568 (Abcam, cat. Number ab175471) was diluted in PBS to a final concentration of 1 μg/mL and incubated for 1h at RT. In order to stain actin, 200 nM SiR actin 674nm (Spirochrome, USA) or Phalloidin iFluor 488 (dilution 1:1000) (Abcam, cat. Number ab176753) were incubated with the secondary antibody solution. In the same solution we also included 1.0 μM Hoechst 33342 (ThemoFisher, cat. Number H3570) or Dapi (Millipore, cat. Number 90229). After 1h the staining solution was removed, and the samples were washed three times with PBS for 10 minutes each step.

For the live/dead assays, the organotypic cell culture systems were incubated 15 minutes before imaging in media with 1 μM calcein AM (Invitrogen) and 1μg/mL propidium iodide (Invitrogen) at 37°C.

### 4.5. Cell imaging

To verify the quality of the printed structures and to observe the cultured cells, an inverted fluorescence microscope IX81 Olympus was used (Plan C objective 4x (NA= 0.1), CAch PhP 10x (NA=0.25) and UCPlan FL N 20x (NA=0.7).

The Nikon Ti2 A1R confocal light scanning microscope located at the Nikon Imaging Center in Heidelberg was used to analyze the actin remodeling after the mechanical stimulations. The objective used for this study was the Nikon Plan Apo λ 20x NA 0.75 (WD 1mm, FOV 0.64 x 0.64mm).

### 4.6. Fish husbandry and maintenance

Medaka fish (*Oryzias latipes*) stocks were kept complying with the local animal welfare standards (Tierschutzgesetz §11, Abs. 1, Nr. 1, husbandry permit AZ35-9185.64/BH Wittbrodt). Fish were maintained as closed stocks in constant recirculating systems at 28°C with a 14h light/10h dark cycle. In this study, a transgenic line carrying a genetic reporter construct marking early retinal cells (Rx3::H2B_GFP) was used[51].

### 4.7. Fish retinal organoids

Retinal organoids were prepared from primary pluripotent embryonic stem cells of Medaka fish (*Oryzias latipes*) as described in Zilova et. al (2021)[50]. In short, embryonic stem cells from blastula stage[52] Medaka embryos were isolated and seeded in Differentiation media (DMEM/F12 (Dulbecco’s Modified Eagle Medium/Nutrient Mixture F-12), GibcoTM Cat:21041025), 5% KSR (GibcoTM Cat: 10828028), 0.1 mM non-essential amino acids, 0.1 mM sodium pyruvate, 0.1 mM β-mercaptoethanol, 20 mM HEPES pH=7.4, 100 U/ml penicillin-streptomycin) into 96 well plates (Nunclon Sphera U-Shaped Bottom Microplate, Thermo Fisher Scientific Cat: 174925). A number of about 150 cells was seeded per aggregate to achieve an organoid size of about 100 µm diameter on day 1 of culture. 15h after seeding, organoids were treated with a final concentration of 2% Matrigel® (Corning, Cat: 356230) for 6h before mounting single aggregates into printed scaffolds.

### 4.8. Printing, stimulation and analysis of Medaka retinal organoid

The procedure to encapsulate the organoid was adapted from a recent publication[53]. Briefly, after printing the structure from IP-PDMS and respective development, the organoid (age D1) was placed manually with the help of a needle inside the structure. Immediately after the mounting, the culture medium was replaced with the HYDROBIO INX N400 (BIO INX) trying to avoid dehydration. The PDMS structure (the outer ring, bridge, and base) was printed at 25x, NA = 0.8 using a commercial laser direct writing device (Photonic Professional GT2, Nanoscribe GmbH). A laser power of 46.2 mW and a printing speed of 60 mm/sec were used as printing parameters. The encapsulation of the retinal organoid was performed with a laser power of 52.8 mW and a printing speed of 70 mm/sec. The mechanical stimulations were performed 24h later using a cantilever with 53.1 N/m stiffness and a tip radius of 24 μm. The load applied was 200 μN with a frequency of 0.5 Hz for 30 minutes. After the stimulation, the organoid was fixed with 4% PFA for 1h. After, the organoid was stained with SiR actin 674nm (Spirochrome, USA) for 1h. Images were acquired with a Nikon Ti2 A1R confocal light scanning microscope equipped with a Nikon Plan Apo λ 20x NA 0.75 (WD 1mm, FOV 0.64 x 0.64mm).

### 4.9. Micro-indentation measurements

To characterize the mechanical properties of the hydrogels where the cells grow on and in, respectively, micro-indentation measurements were performed using a Pavone nanoindentation platform (Optics11 Life) accordingly to a previous published method[54]. The measurements of the HYDROBIO INX N100 (BIO INX) structures were carried out using a cantilever with 0.250 N/m stiffness and a 9 μm tip radius, while for the HYDROBIO INX N400 (BIO INX) we used a cantilever with 0.032 N/m stiffness and a 24 μm tip radius. During the probe calibration and measurements, the structures were kept in cell culture media. After the device was calibrated, indentation control experiments were carried out with 2000 nm indentation depth at a speed of 2 µm/s followed by a hold time of 1 s and subsequent retraction at 2 µm/s on each location. Data analysis was done using the Data Viewer (V2.5.0) software supplied by the device manufacturer. The Young’s modulus E from each load-indentation curve was calculated with the Hertzian contact model using a constant indentation speed as shown by Huth et al.[55]. The contact point of each load-indentation curve was found by using the software integrated contact fit up to 20% of the maximum load. The Hertz fits were applied in the range between contact point (0 nm) and 1400 nm.

### 4.10. Mechanical stimulation of the organotypic cell culture systems

To apply a force to the device and thus mechanically stimulate part of the organotypic cell culture grown in the hydrogel we used Pavone nanoindentation platform (Optics11 Life). During the stimulations the samples were incubated in EBM-2 Medium with standard cell culture conditions at 37°C and 5% CO_2_. The stimulations were performed using a cantilever with 4.10 N/m stiffness and a tip radius of 25 μm. The load applied was 200 μN with a frequency of 0.5 Hz for 30 minutes. During the stimulations we took images in brightfield with a magnification of 20x, and we recorded videos at 24 frames per second. The videos were used for the optical flow analysis in order to determine the displacement of the organotypic cell culture systems.

### 4.11. SEM analysis

Prior to SEM investigations, the IP-PDMS structures were coated with 5nm titanium and 15nm gold respectively using a sputtering system (MED 020, Bal-Tec, Germany). The images were acquired with a Jeol scanning electron microscope (JEOL JSM-7610F) at 5 kV.

### 4.12. FEA analysis

We performed FEA analysis to predict the displacement of the 3D models by using Inventor Autodesk. The displacement was modeled by considering two different materials used to print the structures. In Table 1 (**Supplementary Table 1**) we reported the mechanical parameters used for the analysis. The Young’s Modulus for the IP-PDMS was described in the manufacturers’ datasheet while the shear Modulus was selected from the standard silicone-based materials database. For the HYDROBIO INX N400 we measured the Young’s Modulus from the data reported in **Figure S1**, while for the shear Modulus the value reported in literature for standard gelatin based materials were used[56].

### 4.13. Optical flow analysis

To measure the deformation of the sample, a video (24 frames/sec) of 30 minutes was taken during the indentation process. From the video, image pairs at minimum/maximum strain were extracted for the first 100 oscillations (equal to 3.33 minutes). For each pair, trackable features were chosen from the relaxed image using a Shi-Tomasi corner detector. The displacement for each feature was then determined using Lucas-Kanade optical flow with a window size of 50px (16.5 µm) and pyramid level 2. To remove outliers, the top 5% of displacements, as well as all features outside the structure were excluded. The displacements for all oscillations were then binned into a 20 px (6.6 µm) grid.

### 4.14. Actin alignment characterization

We performed actin alignment characterization by utilizing fast Fourier transform (FFT) of square windows of 51 by 51 pixels chosen from the image in the original space. The windows were overlapped by 50% and an appropriate intensity threshold was considered to filter the blank spaces. The analysis was based on the algorithm developed by S. Marcotti et al.[44] The Python script found in https://github.com/OakesLab/AFT-Alignment_by_Fourier_Transform/tree/master/Python_implementation has been modified for this purpose.

## Supporting Information

Supporting Information is available from the author.

## Acknowledgements

The work was supported by the Deutsche Forschungsgemeinschaft (DFG, German Research Foundation) under Germany’s Excellence Strategy via the Excellence Cluster 3D Matter Made to Order (EXC-2082/1–390761711) (C. S. U, J. W.) and the Carl Zeiss Foundation through the Carl-Zeiss-Foundation-Focus@HEiKA (C. S. U.). F. C. particularly thanks 3DMM2O for the “PostDoc Take-Off Grant 2022”.

F.T. acknowledges support by the Deutsche Forschungsgemeinschaft (DFG; German Research Foundation) through SPP SE 1801/5-1, (2303731). This work was funded in part by the Volkswagen Foundation through the Initiative “Life?,” Az. 96733, the Max Planck School Matter to Life supported by the German Federal Ministry of Education and Research (BMBF) in collaboration with the Max Planck Society. C. S. U., M. V. and F. C. also thank the European Research Council for support through the Consolidator Grant PHOTOMECH (Grant no. 101001797). The authors thank the Nikon Imaging Center in Heidelberg and the Soft (bio)materials characterization Core Facility (Biomechanics) at IMSEAM (Heidelberg University) for their support.

## Supplementary Video

**Video S1.** Video acquired during the hydrogel printing process.

**Video S2.** Video of the mechanical stimulation. The duration (15 seconds) and speed (2x) of the video is adapted to reduce the file size.

**Video S3.** Video acquired during the mechanical stimulation of the Medaka retinal organoid.

## Table of content (Toc)

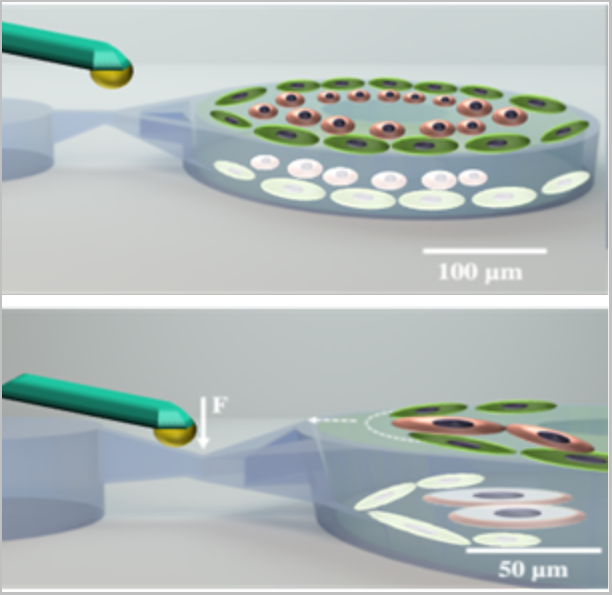

We have developed a multimaterial 3D structure printed via two-photon polymerization that can be mechanically stimulated with micrometer-scale precision by a cantilever. The inner cylindrical part of the structure is printed with a hydrogel that allows direct printing of the cells forming an organotypic cell culture. Its stimulation causes remodeling of the cytoskeleton and cell morphology.

---

**Figure S1.**
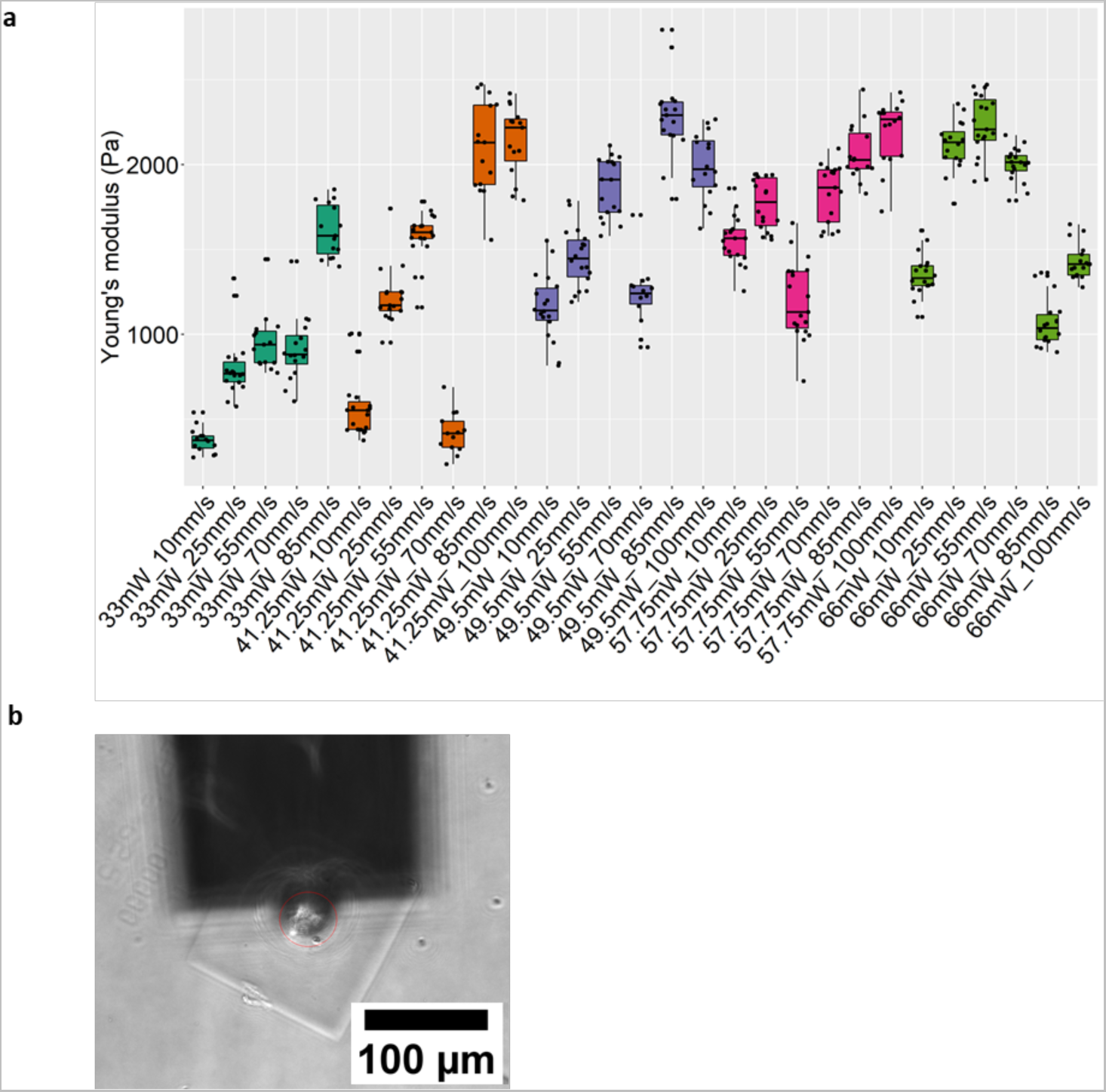
Characterization of the hydrogel stiffness for direct cell printing. a) Boxplot of the Young’s Modulus (Pa) of different cubes printed with HYDROBIO INX N400 ink with increasing laser power and speed. c) Representative image acquired during the analysis with the nanoindenter.

**Figure S2.**
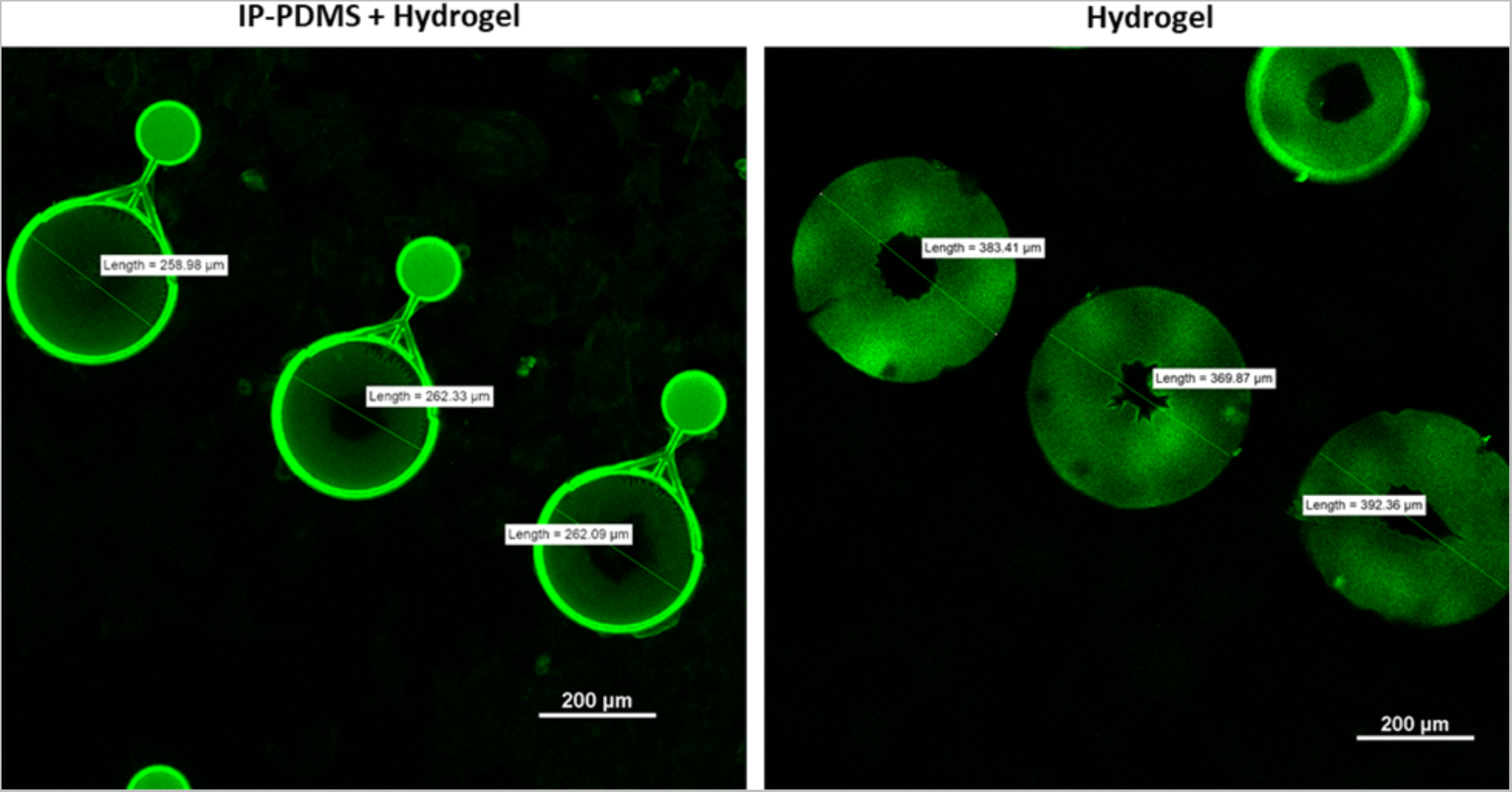
Swelling of hydrogel cylinders. The two pictures compare cylinders of hydrogel (HYDROBIO INX N100) with and without the IP-PDMS structures. Those without IP-PDMS swell to an average diameter of 380 μm.

**Figure S3.**
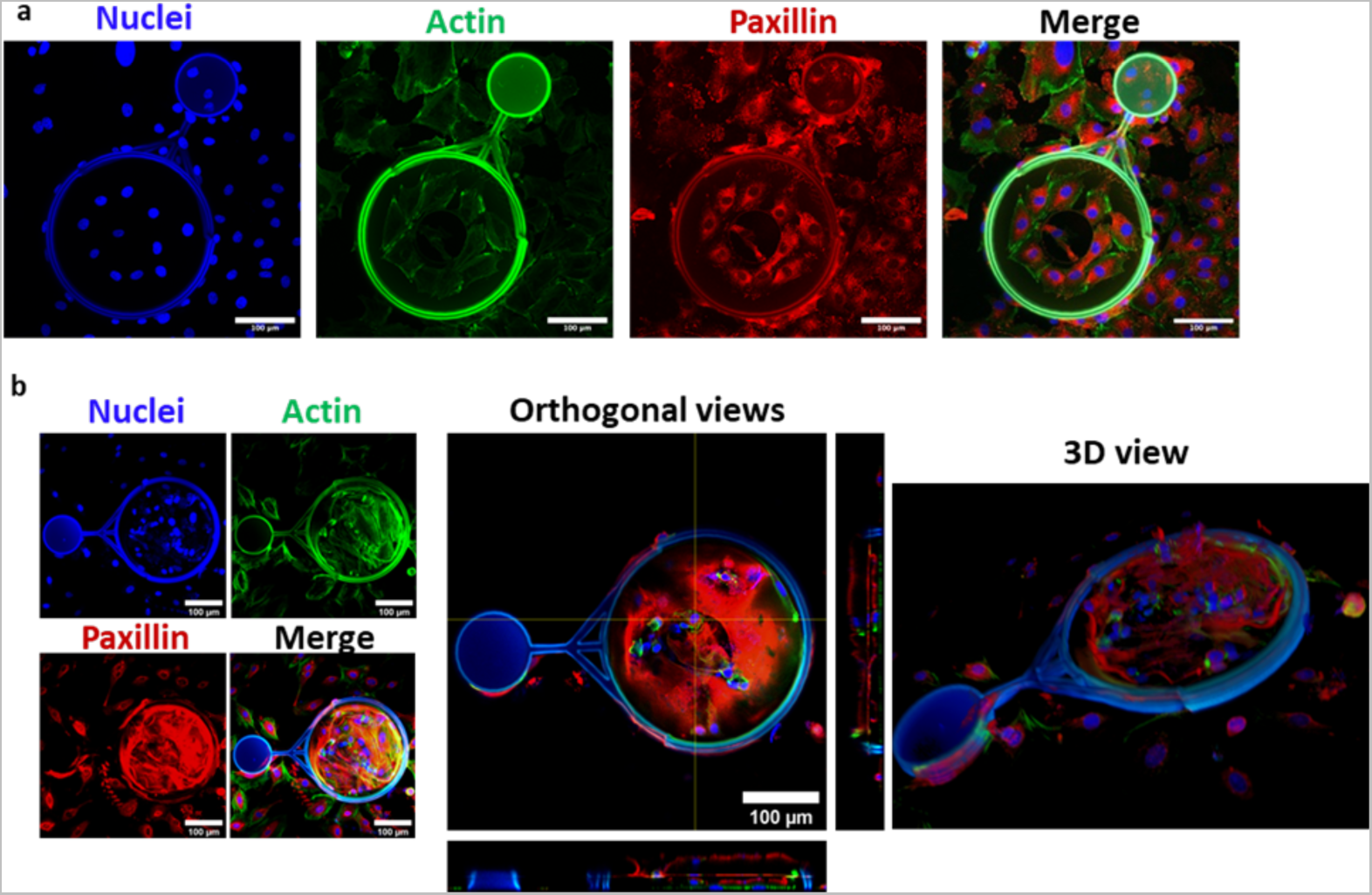
Incubation of REF cells on top of the 3D structures. a) REF cells (7.500 cells/coverslip) were incubated for 4 days with the structures and then stained for Actin (green), Paxillin (red) and Nuclei (Hoechst, blue). Scale bars 100μm. b) Confocal images of the structures incubated with 50.000 cells/coverslip. As for the previous images, the structures were stained for Actin (green), Paxillin (Red) and Nuclei (Hoechst, blue). In the middle we showed the orthogonal view to highlight the presence of the cells in the Z-axis. On the right side of the figure we showed the 3D view. Scale bars 100μm.

**Figure S4.**
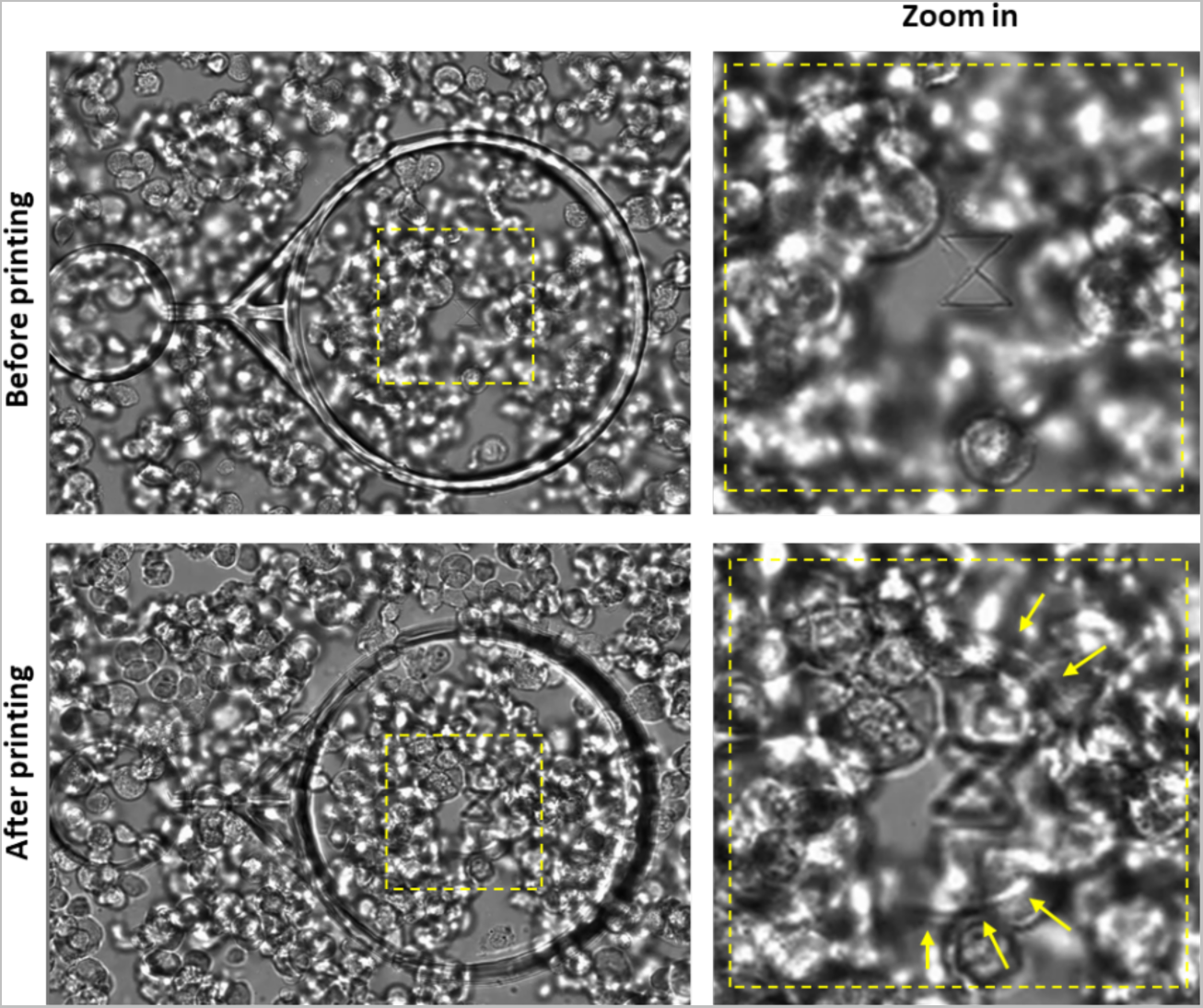
Direct printing of cells into the hydrogel. The image above is extracted from the **Video S1** (refer to the video supplement section) prior to starting the direct printing of cells mixed within the hydrogel. Below is the picture taken after completing the printing process. In the right column, we have included a zoomed view highlighting the edge of the hydrogel, formed at the end of printing, using yellow arrows.

**Figure S5.**
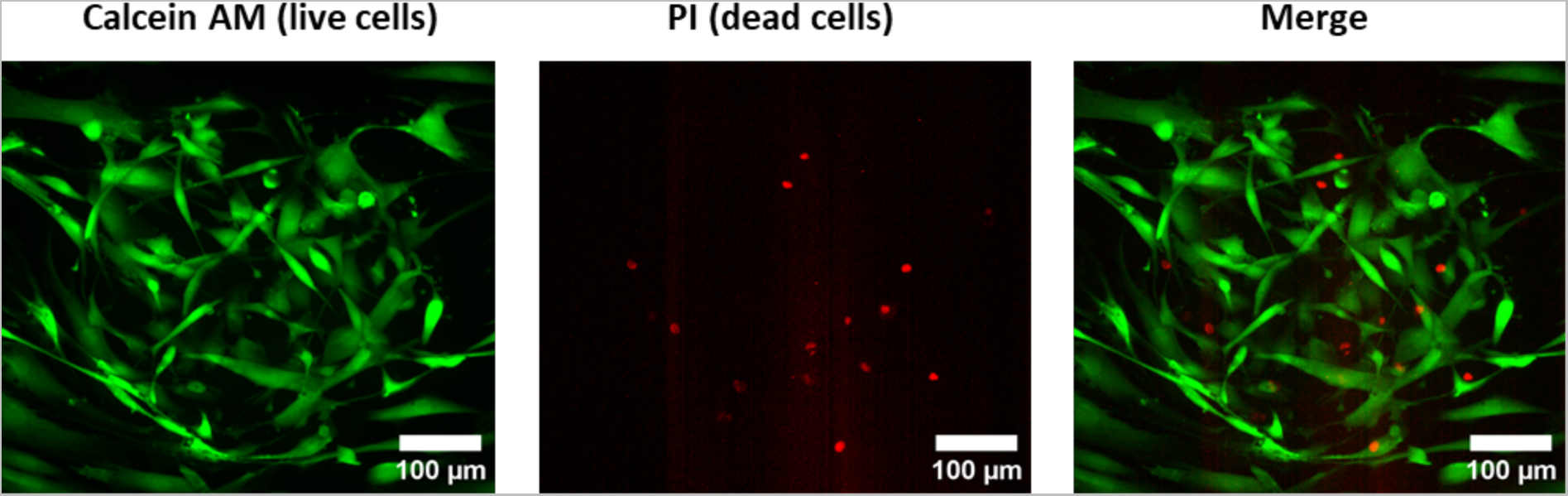
Live/Dead assay. HFLS directly printed in the hydrogel (HYDROBIO INX N400) without the IP-PDMS structure were stained with calcein AM (green) and propidium iodide (red) 24h after printing.

**Figure S6.**
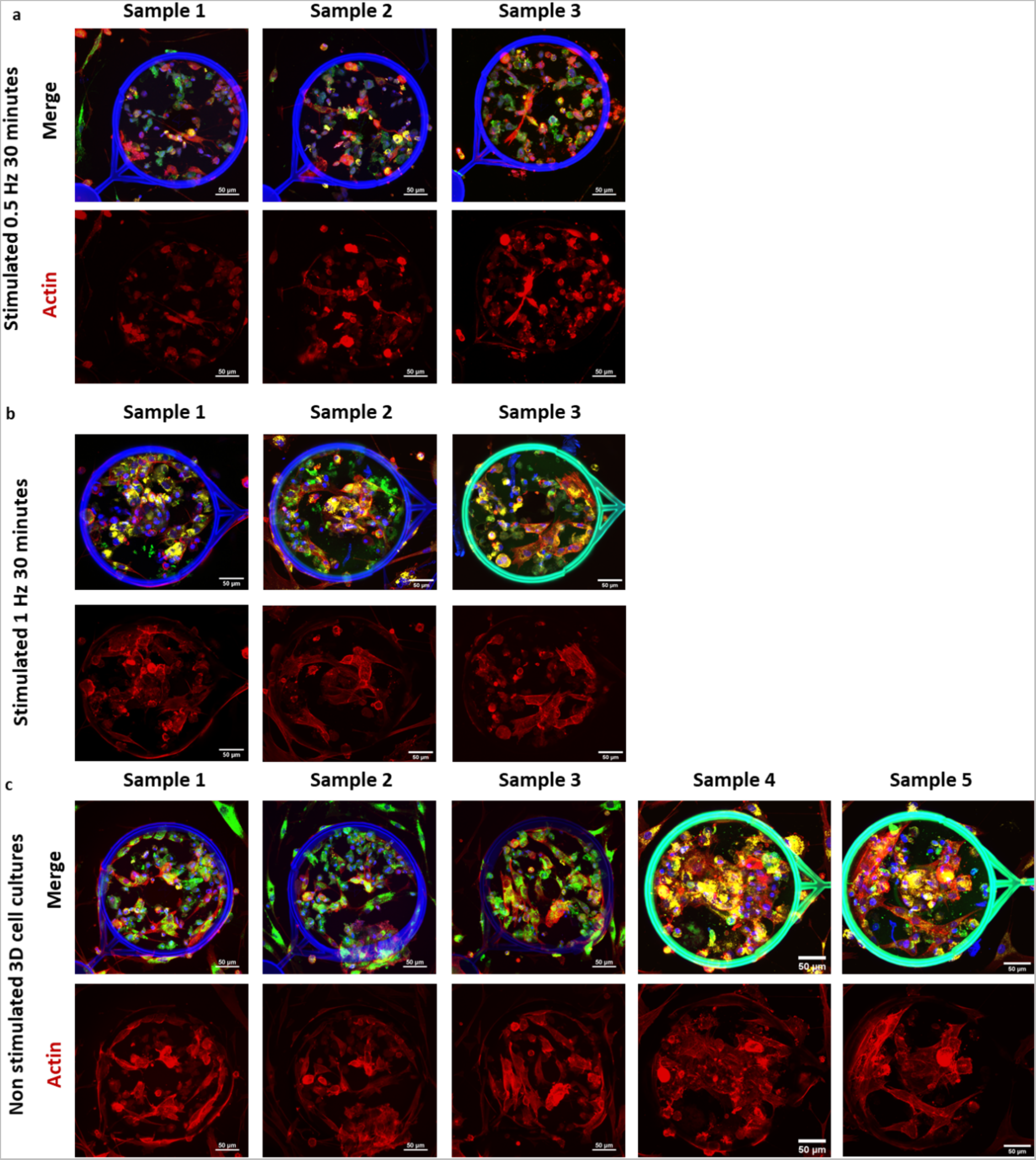
Actin remodeling in mechanically stimulated organotypic cell culture systems. a) 3D organotypic cell culture systems mechanically stimulated for 30 minutes with a load of 200 μN at 0.5Hz. In the Top row: merge of the max intensity projections of 3 channels: HFLS cells (green), Huvec cells (yellow), Nuclei (Hoechst). b) 3D organotypic cell culture systems mechanically stimulated for 30 minutes with a load of 200 μN at 1 Hz. c) Non-stimulated 3D organotypic cell culture systems. Images were colored according to the scheme above. Scale bars 50 μm.

**Table S1.** Mechanical properties of the materials used for the simulations.

|  |  |  |  |
| --- | --- | --- | --- |
| Name | IP-PDMS |  | HYDROBIO INX N400 |
| Stress | Young's Modulus | 0,015 GPa | 0,000004986 GPa |
|  | Poisson's Ratio | 0,49 | 0,49 |
|  | Shear Modulus | 0,00503356 GPa | 0,00000167315 GPa |

